# Genome-wide signatures of local adaptation among seven stoneflies species along a nationwide latitudinal gradient in Japan

**DOI:** 10.1101/440552

**Authors:** Maribet Gamboa, Kozo Watanabe

**Author notes:** Correspondence: Maribet Gamboa.

## Abstract

**Background:** Environmental heterogeneity continuously produces a selective pressure that results in genomic variation among organisms; understanding this relationship remains a challenge in evolutionary biology. Here, we evaluated the degree of genome-environmental association of seven stonefly species across a wide geographic area in Japan and additionally identified putative environmental drivers and their effect on co-existing multiple stonefly species. Double-digest restriction-associated DNA (ddRAD) libraries were independently sequenced for 219 individuals from 23 sites across four geographical regions along a nationwide latitudinal gradient in Japan.

**Results:** A total of 4,251 candidate single nucleotide polymorphisms (SNPs) strongly associated with local adaptation were discovered using Latent mixed models; of these, 294 SNPs showed strong correlation with environmental variables, specifically precipitation and altitude, using distance-based redundancy analysis. Genome–genome comparison among the seven species revealed a high sequence similarity of candidate SNPs within a geographical region, suggesting the occurrence of a parallel evolution process.

**Conclusions:** Our results revealed genomic signatures of local adaptation and their influence on multiple, co-occurring species. These results can be potentially applied for future studies on river management and climatic stressor impacts.

## Background

Understanding the influence of environment on genomic variation in natural populations is the primary goal for addressing questions regarding evolution. Principally, divergence selection derived from natural selection may result in allele frequency shifts at loci under selective pressures, such as heterogeneous climatic factors, to maximize fitness in local environment (1). Therefore, the capacities of organisms to respond to changing selective pressures (such as climate) may widely vary due to their potential pending genetic adaptation and the rate at which new genomic variations arise (2, 3).

Studies investigating the genomic basis of adaptation often compare genomic information of populations along heterogeneous environmental (2). First, the genome was screened for outliner loci by determining associations between allele distributions and environmental variables that were presumed to be selection drivers (4, 5, 6). Then, correlations among environmental variables combined with population genomic approaches (i.e., fixation index and inbreeding coefficient) may complement each other in terms of power and accuracy (7). However, discerning the relative contribution of environmental conditions in shaping genetic diversity remains a challenge. Identifying factors such as natural selection, genetic drift, and dispersion that modify patterns of genomic variation remains difficult (8).

Genome–environmental association analysis along environmental gradients (9) provides an opportunity to examine the influence of environmental conditions in shaping genetic variation of natural populations. Environmental information, as a combination of landscape effects (10) and environmental clines (11), provides stronger evidence for examining adaptive divergence. This type of analysis considers the effect of population structure and geographical scale and expects that genome signatures of selection become more pronounced with increasing geographical scale owing to larger environmental gradient differences (12). Genetic response to environmental gradients has been evidenced in a wide range of taxa, including plants (13, 14), fish (15), and invertebrates (16, 17). Overall, reported studies have highlighted that the selective pressure resulting from environmental changes can be explained by the evolutionary response of population along these gradients. However, these studies investigating this association have been based on a single species (14, 15, 18, 19), thus neglecting the potential effect of this association on multiple species that co-occur in the same habitat.

The occurrence of multiple species in a community remains another challenging aspect of determining the influence of environment on genomic variation. Adaptive genomic variation is strongly influenced by the complex dynamics between environmental and community effects (20). Due to evolutionary dynamics, the assessment of genomic variation in organisms that co-occur with other species may not match that predicted using single species approaches (21) because of the effect of competition, predation, and co-evolution. For example, using multiple-species observations, it is possible to observe genome–genome interactions (e.g., hybridization and mutualistic), genome–environment interactions (e.g., parallel adaptation, recombination rate, and demography), and genome-genome-environment interactions (e.g., interactions among species) (20) between organisms. Data on the impact of environmental change on evolutionary processes at the community and ecosystem scale is lacking (20). In order to gain a better understanding of adaptive genomic variation, genetic consequences of co-existing species in a local community should be studied (22).

Aquatic insects are ideal organisms for observing genome–environment interactions. Stoneflies (Plecoptera) are aquatic insects that exhibit limited airborne dispersal within stream corridors (23). Stoneflies are considerably more sensitive to environmental changes, such as low oxygen concentration and high water temperature, than other aquatic insects (24). Until now, studies evaluating genome–environmental association in aquatic insects have often used mitochondrial DNA sequences with a particular molecular marker (Sanger sequencing) (25), microsatellite (26), or Amplified Fragment Length Polymorphism makers (27); however, NGS data has been scarcely used except for in few studies such as those on damselfly (16) and chironomidae (17).

In this study, we examined the genome-wide signatures of adaptive divergences in seven species of stoneflies using a double-digest restriction-site-associated DNA protocol (ddRAD). Several techniques have been applied for the detection of adaptive genes, such as direct sequencing of genomic regions of interest (28), genome re-sequencing (18), genotyping by sequencing (GBS; 14), transcriptomics (19), and restriction site associated DNA sequencing (RAD-Seq; 15). Among these techniques, RAD-Seq allows the analysis of multiple organisms at the same time (29) and the genomic analysis of organisms without a reference genome (30). In addition, it allows for the identification of thousands of genetic markers randomly distributed across the genome (31), which further increases the chances of detecting loci under natural selection. Therefore, RAD-Seq has been employed for understanding the pattern of genetic structure of populations across environmental heterogeneity (e.g. 32).

Here, we performed population and landscape genomic studies using ddRAD so as to screen imprints of selection on a national-wide scale in Japan archipelago. We primarily aimed to evaluate the relative strength of selection in influencing the distribution of genomic variation at different spatial scales, to identify putative environmental drivers, and to observe their effects on the community. For this, we collected samples along a nationwide latitudinal gradient in Japan with varying meteorological and hydrological conditions. We applied an individual-based Latent Factor Mixed Model (LFMM) and a multivariate redundancy analysis (RDA) to identify putative single nucleotide polymorphisms (SNPs) under selection and the main environmental population differentiation driver. The resulting species-specific association of SNPs with environmental variables were compared to determinate the degree of nucleotide changes (i.e., nucleotide composition) among the seven species so as to possibly explain their local adaptation. Although the use of candidate SNPs has been the standard method to determine loci putatively under selection, integrating environmental gradients and multiple species can be reliably used to identify such loci and selective agents underlying the observed patterns either in a population or community.

## Methods

### Sample collection

We ad hoc collected a total of 47 species. Of these, eight species that commonly occurred throughout four regions in Japan were shortlisted to test the potential effect of climatic variation in these regions. The eight species of stream stoneflies selected (*Perlodini incertae*, *Haploperla japonica*, *Nemoura ovocercia*, *Rhabdiopteryx japonica*, *Obipteryx femoralis*, *Isoperla nipponica*, *Amphinemura longispina*, and *Stavsolus japonicus*) have different biological requirements, such as feeding behavior, and habitat preferences. An average of 31 stonefly nymphs per species were collected in January–March 2014 along a latitude cline in Japan using D-flame nets (mesh size = 250μm). Collected specimens were identified using the taxonomic key of Japanese aquatic insects (33).

Sampling of stonefly nymphs was carried out at 23 sites across four regions with different climatic conditions, including: Matsuyama (6 sites), Gifu (4 sites), Sendai (6 sites), and Sapporo (7 sites) (Fig. 1, Additional file 1: Table S1, Additional file 2: Table S2). Sampling sites in Matsuyama located in Ehime Prefecture in Shikoku Island, South of Japan (geographical distance to other regions ranged 404–1295 km) has a mean temperature of 17°C and an altitude range of 154–277 meters above sea level (masl). The Gifu region, situated in Gifu prefecture at the central area of Japan (geographical distance to other regions range 374–959 km) has a mean temperature of 15°C and an altitude range of 251–720 masl. The Sendai region, located in Miyagi prefecture in the northeast area of Japan (geographical distance to other regions range 374–895 km) has a mean temperature of 12°C and an altitude range of 261–394 masl. The sampling sites in the Sapporo region, in Hokkaido prefecture positioned at the Northern most part of Japan (geographical distance to other regions range 418–1295 km) has a mean temperature of 7°C and an altitude range of 95–298 masl.

**Figure 1.**
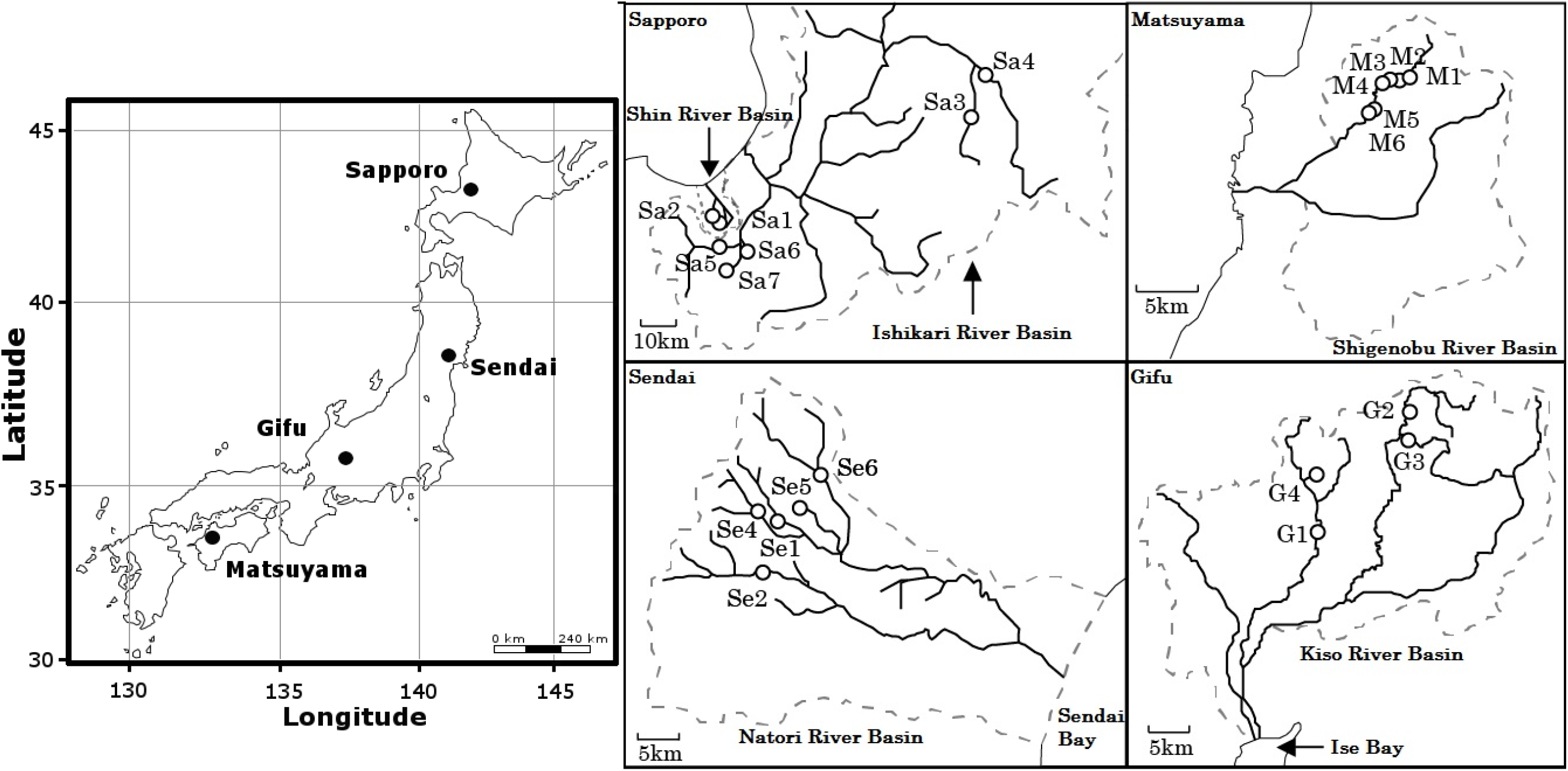
Map depicting areas of Japan used for sampling stonefly nymphs for this study. Filled circles indicate the four geographical regions (left) included in the study, and open circles indicate the sampling sites within each region (right)

Data on precipitation (mm), river water level (m), discharge (m^3^/s), water temperature (°C), air temperature (°C), and depth of snow cover (cm) for the sampling sites (Additional file 1: Table S1) were obtained from the Ministry of Land, Infrastructure, Transport and Tourism of Japan (http://www1.river.go.jp/) and the NASA Giovanni website (https://giovanni.sci.gsfc.nasa.gov/giovanni/) within standard NASA estimate algorithms. Environmental data for each sampling site was collected from a different meteorological station. Different meteorological stations were situated within 2.5–19.8, 7–52, 1.3–16, and 17–103 km in Matsuyama, Gifu, Sendai, and Sapporo regions, respectively.

Geographic distances between sampling sites were calculated from the transformation of GPS coordinates (34) using great-circle distances calculated for the Vincenty Ellipsoid in geosphere package of R v.3.3 (35) and used for principal components analysis (PCA) using the stats package in R v.3.3 (R Core Team, www.r-project.org). The results obtained from PCA were used to statistically separate sampling sites.

### ddRAD library preparation and sequencing

A total of 270 stoneflies belonging to eight species (average = 31 individuals per species; range = 20–52 individuals per species) were used for RAD-Seq. All individuals were washed thrice with ethanol to eliminate any contamination. Genomic DNA was extracted from each individual using DNeasy tissue kit (Qiagen GmbH, Hilden, Germany) following manufacturer’s instructions. This DNA extraction method involved the use of the whole body without any destruction of the specimen, thus reducing the probability of contamination (gut content contamination). ddRAD libraries were constructed as previously described (29) with slight modifications. Briefly, 100 ng genomic DNA per sample was digested with restriction endonucleases, *Sbf*I (NEB) and *Hae*III (Takara) at 37°C for 3 h. Restriction fragments of each sample were ligated to a unique 6-mer adaptor. Ninety samples with equimolar DNA concentrations were combined and electrophoresed on agarose gels. Subsequently, 300–500 bp fragments were excised and purified using MinElute Gel Extraction Kit (Qiagen).

Purified DNA samples were PCR amplified using two sequential PCRs. The first PCR was carried out in a 20 μl reaction, consisting of 12 μl of Phusion High-Fidelity PCR Master Mix with HF buffer (NEB), 2 μl DNA template, 2 μl of each forward and reverse primers specific to Illumina adaptors, 1 μl of Betaine (Sigma), 1 μl of DMSO (NEB), and 2 μl of water, using the following conditions: initial denaturation at 98°C for 1 m, followed by 18 cycles of 98°C for 10 s, 62°C for 30 s and 72°C for 30 s, and a final extension at 72°C for 10 m. The second PCR was carried out to eliminate non-specific sequences using 4 μl of Phusion HF buffer (NEB), 2 μl of dNTPs (Promega), and 2 μl of each forward and reverse Illumina sequencing primers, using the following conditions: 98°C for 3 min, 60°C for 2 min, and 72°C for 12 min.

Five PCR replicates were pooled to form a single library. Each library was quantified using KAPA library quantification kit (Roche). Three DNA libraries each comprising 90 individuals were sequenced per lane using HiSeq 2500 Illumina sequencer (paired end, 2 x 100 bp) at the Beijing Genomics Institute, China. One species (*Perlodini incertae*, 51 individuals) was removed from the analysis because of its low read recovery (<8,000,000 total reads). Therefore, we used seven species (219 individuals) in the analysis.

### Data processing

Forward and reverse raw data (789,587,765 reads) were screened for tags and adapters using CUTADAPT (36) (Additional file 9: Figure S1). A phred-type quality score of Q20 was used to trim sections of the reads using FASTX toolkit v.0.0.13 (http://hannonlab.cshl.edu/fastx_toolkit/). Reads shorter than 50 bp were excluded. The number of reads was reduced to 675,870,356 (86% of the total reads).

RAD sequences were processed using a pipeline in STACKS v 2.0 that incorporates a maximum likelihood statistical model to identify sequence polymorphisms (37). Sequences were demultiplexed and quality filtered using the process_RADtags program. Reads shorter than 50 bp with a quality read score of < 20 or with ambiguous barcodes (i.e. erroneous barcodes) were removed. The number of reads was reduced to 675,100,758 (85% of the total reads).

*De novo* assembly of short paired-end reads was performed independently using three programs in the STACKS software package. The ustacks program uses a set of reads as input and aligns them into exactly matching putative alleles. Different parameter combinations were evaluated (Additional file 3: Appendix S1, Additional file 4: Table S3, Additional file 5: Table S4). The analysis revealed similar genomic comparisons [i.e. observed and expected homozygotes and heterozygotes, fixation index (*F*_st_, Additional file 6: Table S5), and inbreeding coefficient (*F*_is_)] but different number of loci (Additional file 5: Table S4). For the following analysis, parameter combinations with a high number of polymorphic loci with a depth read coverage ranging from 30X to 60X were used. We applied the following conditions as previously recommended by (38, 39): minimum read depth to create a putative allele group = 3, number of mismatches allowed between putative alleles within individuals = 2, and number of mismatches allowed between putative alleles within a putative allele group = 2. Reads that failed to meet these criteria were removed. This process reduced the number of reads to 499,574,561 (63% of the total reads).

The output file from ustacks was a catalog of putative alleles per individual. This catalog was used as an input file for the cstacks program implemented in STACKS. This program merges putative alleles together and creates a set of consensus putative loci. The default parameters, namely permitting mismatches between putative alleles of two nucleotides and removing putative alleles considered unique alleles were applied. This process reduced the number of reads to 310,722,904 (41% of the total reads).

The output list of consensus putative loci was used as an input file for the sstacks program implemented in STACKS. This program matches each individual putative locus list against the consensus putative loci for each species. The default parameters were applied, following the base matching on the alignment position, which allowed the detection of sequences errors. Consensus putative loci for each species that were found among all individuals were retained. The tsv2 and gstacks programs were implemented in STACKS to assemble and merge paired-end reads. These processes reduced the number of reads to 100,980,315 (16% of the total reads) among 50,514 loci (Additional file 9: Figure S1).

Following *de novo* assembly, we performed an additional data-filtering step using the populations program implemented in STACKS. Consensus putative loci were excluded from the study using the following criteria: 1) present in <80% of individuals; 2) allele frequency < 0.05; 3) <30X – >60X read coverage; and 4) > one SNP per locus. We examined whether a high read depth affected our data in the filtering as well as the downstream analysis (additional file 3: Appendix S1) as previously reported (40). A cut-off filtering process was employed using vfilter by examining whether the high depth coverage reads affected our data (https://github.com/bcbio/bcbio-nextgen) following a previous study’s suggestion (40) (the output removed 0.001% of the total reads). The number of reads was reduced to 64,502,837 (8% of the total reads, Additional file 7: Table S6) among 24,318 loci (Additional file 8: Table S7, Additional file 9: Figure S1).

The observed (*H*_o_) and expected heterozygosity (*H*_e_), *F*_is_, and *F*_st_ were estimated per region and per species using popGenome package in R v3.3 (41). The significance of pairwise *F*_st_ values was tested using 10,000 permutations.

Population genetic structure was explored using fastSTRUCTURE (42) without *a priori* information of the geographical origin of each sample. Analyses were performed using the admixture model with correlated allele frequencies and a burn-in period of 200,000 MCMC interactions, followed by 300,000 interactions per run using the logistic parameter. We selected a number of K (putative populations) ranging from 1 to 12 with all loci. Five replicate analyses were performed for each putative population (K). The number of cluster was inferred using the complexity model of fastSTRUCTURE. The results of 5 replicates of the selected K values were pooled into a single result using CLUMPAK web version (http://clumpak.tau.ac.il/index.html) and their consistency was examined following (43) recommendations using CLUMPP V. 1.1.2 (44) by the Greedy algorithm.

### Power analysis

To determine the power of RAD datasets to resolve population structure, we performed a power analysis using POWSIM v. 4.1 (45). This program estimates the statistical power of genetic homogeneity in individual species. To reflect our sampling design, we set the number of subpopulations to four, with 5–10 samples per subpopulation for SNPs (based on the distribution of the individuals per sampling site; Additional file 2: Table S2). We set the effective population size of the subpopulations to 1,000, 2,000, and 3,000, and we adjusted the generation time (t) to assess power at multiple F_ST_ values (10 and 20 generations) as previously described (46). F_ST_ in Powsim assumes the independence of the subpopulations (45). Power was expressed as the proportion of significant outcomes for 1,000 replicates and a statistically significant test (p < 0.05).

Additionally, the power of our SNP data to the environmental association was determined using Quanto v.1.2.4 (47). We adjusted the parameters based on our sample size (Additional file 6: Table S5) and implemented independent individuals for the gene-environment interaction, a marginal effect, a population mean of 20 per region, and a marginal R squared model.

### Genome-environmental associations

The latent factor mixed model (LFMM; 48) was used to identify local adaptations among stonefly populations. LFMM detects correlations between environmental variables and genetic variation through the estimation of latent factors and regression coefficient. For LFMM analysis, data on SNPs, population structure, and environmental variables and spatial eigenvectors among sampling sites [8–22 (mean, 14.4) sites per species; Additional file 2: Table S2) were used with the LEA package in R v.3.3 (49). To eliminate false positives, Benjamin Hochberg algorithm was applied using the q value package in R v.3.3 (50) at a false discovery rate (FDR) of 10% following the manual of LEA package (49).

Multidimensional scaling (MDS) of all candidate SNPs generated using LFMM and their correlation with environmental variables was examined to determine the proportion of genomic variation affected by environment. Candidate SNPs were extracted using VCFtools v.0.1.14 (51) and used to calculate the allele frequencies (identity by state; IBS) at each site using Hamming distance matrix with PLINK v.1.9 (52). The IBS matrix was used for Principal Coordinate Analysis (PCoA). The resulting eigenvalues of PCoA were used for the subsequent distance-based redundancy analysis (db-RDA; 53). Statistical significance of db-RDA models was tested using ANOVA. The db-RDAs were conducted using capscale and ordistep functions and by plotting MDS1 and MDS2 in the vegan package in R v3.0.2 (R Core Team; 54). MDS analysis was also used to enable the identification of a SNP associated with the respective environmental variable; therefore, we examined correlations between environmental variables and individual candidate SNPs using MDS and db-RDA.

### Adaptive genetic variation among species

To determine the degree of adaptive genetic variation in stonefly genomes, sequences of loci of seven stonefly species harboring candidate SNPs were aligned using bowtie2 v.2.3 (55). Default parameters were used in end-to-end alignment that allowed up to two mismatches per 90 base reads. Subsequently, the best match sequences for each locus were identified. Additionally, these sequences under putative selection were used to compute mean nucleotide substitution rate within species (i.e., individuals within species) and within or between geographical regions (i.e., species per region) by pooling the seven species (41). Mean nucleotide substitution rate was calculated using pairwise nucleotide substitution rate, where the expected value ranged from 0 (similar) to 1 (dissimilar).

We identified the gene function of the sequences of the candidate loci using web version of tBLASTx against the nucleotide databases of GenBank, EMBL, DDBJ, PDB, RefSeq, PDB, SwissProt, PIR, and PR in NCBI (http://blast.ncbi.nlm.nih.gov) under default parameters. Sequences with an e-value ≤1e-5 and ≥90% sequence similarity were used in Gene Ontology analysis using web service Uniprot database (https://www.uniprot.org). We excluded a possible contamination and an erroneous gene function in our sequences by processing our DNA sequences against the entire NCBI database using DeconSeq web interface (http://deconseq.sourceforge.net/).

## Results

### Data filtering

RAD-Seq generated a total of 789,587,765 raw reads with an average of 263,195,922 reads per library, 1,802,711 reads per individual, and 3,527,060 per species per region. The length of raw reads ranged from 90–100 bp with an average length of 98 bp. In total, 113,717,409 (14%) reads with low quality, ambiguous barcode and short length were discarded. On an average, 3,606,439 reads per individual (range: 2,600,058–4,016,315 reads) with an average length of 98 bp (range = 90–100 bp) were successfully aligned generating 53,835 consensus loci (supplementary Fig. 1). After loci identification and SNP calling, a dataset of 50,154 loci representing 16% of the total reads was generated, which was equivalent to 222–4,359 loci per species (Additional file 9: Figure S1). After final data filtering (i.e., removing consensus putative loci that were present in less than 80% of individuals), 24,318 SNP loci were identified with 202-2,167 SNP loci per species (mean = 868 SNP loci per species, Additional file 8: Table S7) and an average 45-fold coverage per species (range = 30–60-fold).

The statistical power of our SNP dataset to population structure (POWSIM program) and to gene-environmental association (Quanto program) was evaluated. Both programs provided consistently high power to detect genetic differences (90% mean statistical significance) and detection of association (57% average chance of detection) among the populations.

### Population structure

Population statistics are summarized in Table 1. Fixation index (*F*_st_) and inbreeding coefficient (*F*_is_) within species varied between 0.043–0.123 and 0–0.032, respectively. Expected heterogeneity (He) was an average of 0.33 for all species without spatial differentiation between regions.

**Table 1.** Summary of sample size and molecular metrics. Where: *n*, number of individuals; *H*_o_, average observed heterozygosity; *H*_e_, average expected heteozygosity; *F*_st_, mean genetic divergence of average pair geographical locality; *F*_is_, inbreeding coefficient

| Species | $n$ | Total reads | Total loci | $H_o$ | $H_e$ | $F_{st}$ | $F_{is}$ |
| --- | --- | --- | --- | --- | --- | --- | --- |
| *Nemoura ovocercia* | 20 | 2532235 | 10448 | 0.71 | 0.3 | 0.106 | 0.032 |
| *Haploperla japonica* | 32 | 495817 | 1624 | 0.75 | 0.20 | 0.078 | 0 |
| *Stavsolus japonicus* | 52 | 798464 | 7018 | 0.77 | 0.22 | 0.043 | 0.027 |
| *Rhabdiopteryx japonica* | 22 | 2320398 | 9248 | 0.75 | 0.27 | 0.093 | 0.014 |
| *Obipteryx femoralis* | 31 | 3318488 | 12411 | 0.80 | 0.20 | 0.069 | 0.011 |
| *Isoperla nipponica* | 32 | 942014 | 3086 | 0.70 | 0.29 | 0.081 | 0.003 |
| *Amphinemura longispina* | 30 | 572899 | 6319 | 0.70 | 0.26 | 0.120 | 0.050 |

Bayesian clustering analysis revealed 3 to 6 populations (K) as the most parsimonious partitioning of individuals within species. Whereas *K* = 3 in *H. japonica* [ln(*K*) = −0.079] and *S. japonicus* [ln(*K*) = −0.040]; *K* = 4 in *A. longispina* [ln(*K*)= −0.044], *O. femoralis* [ln(*K*)= 0.038], *R. japonica* [ln(*k*)= 0.013], and *N. ovocercia* [ln(*K*)= −0.003]; and *K* = 6 in *I. nipponica* [ln(*K*) = 0.008). Random spatial distribution of populations between regions was detected; of the four populations identified in *N. ovocercia*, one population was distributed across 3 of 4 sampling regions, except Matsuyama.

### Genome-environmental association

Of the 24,318 SNPs, 4,251 (18%) candidate SNPs (148–1,910 SNPs per species) identified by LFMM were potentially associated with regional environment (Fig. 2, Table 2). The highest number of SNPs associated with environmental variables was identified in *R. japonica* (1,910 candidate SNPs), followed by *N. ovocercia* (922 candidate SNPs).

**Figure 2.**
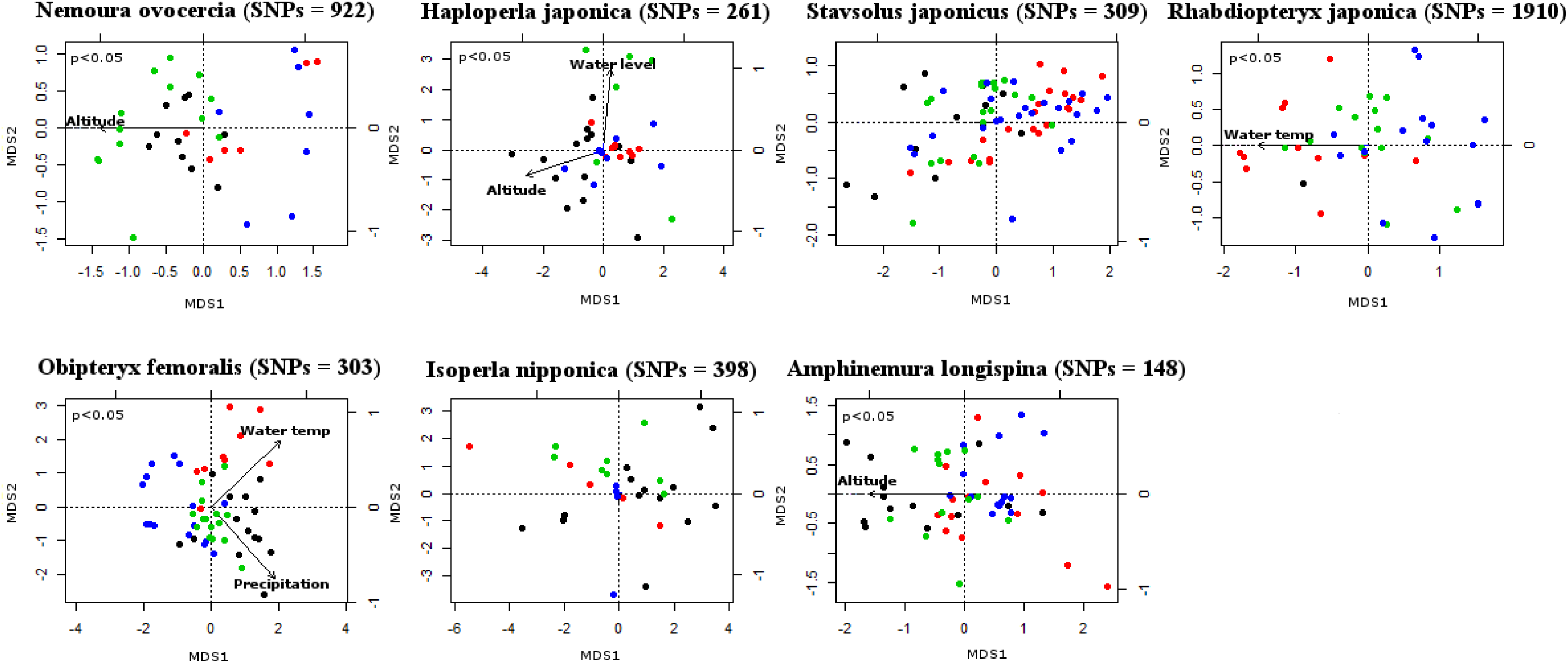
Genome-wide multivariate genome-environmental associations. Panels shows the multidimensional scaling analysis (MDS1 and MDS2) of 4,251 candidate SNPs of seven stoneflies species across four regions of Japan. Colored circles represent different regions: red, Matsuyama; black, Gifu; green, Sendai; and blue, Sapporo

**Table 2.** db-RDA analysis of SNPs and environmental variables for each species. Where: H, number of haplotypes; k, number of populations; spatial and environmental correlation: a, altitude; p, precipitation (mm); wl, water level (m); d, discharge (m^3^/s); sc, snow cover (cm); wt, water temperature (°C);

| Species | SNPs | H | k | Candidate | Unique | Correlated | Spatial and environmental correlation |  |  |  |  |  |  |
| --- | --- | --- | --- | --- | --- | --- | --- | --- | --- | --- | --- | --- | --- |
|  |  |  |  | SNPs | SNPs | SNPs | a | p | wl | d | sc | wt | at |
| *Nemoura ovocercia* | 5073 | 51230 | 4 | 922 | 30 | 66 | 24 | 5 | 5 | 11 | 5 | 5 | 11 |
| *Haploperla japonica* | 943 | 5987 | 3 | 261 | 8 | 6 | 3 |  | 3 |  |  |  |  |
| *Stavsolus japonicus* | 3283 | 58180 | 3 | 309 | 16 | 10 | 2 | 6 |  | 2 |  |  |  |
| *Rhabdiopteryx japonica* | 4572 | 71510 | 4 | 1910 | 113 | 52 | 4 | 12 | 4 | 4 | 12 | 12 | 4 |
| *Obipteryx femoralis* | 6445 | 18320 | 4 | 303 | 8 | 24 | 4 | 5 | 3 |  | 3 | 5 | 4 |
| *Isoperla nipponica* | 1434 | 24405 | 6 | 398 | 8 | 92 |  | 31 | 31 |  |  |  | 30 |
| *Amphinemura longispina* | 2568 | 17512 | 4 | 148 | 22 | 44 | 10 | 6 |  |  | 10 | 8 | 10 |

Multidimensional scaling (MDS) of all candidate SNPs showed genomic variation affected by environment (Fig. 2). The average amount of genotypic variation explained by the first two MDS axes was 76% for seven species. MDS2 (overall species average of 33% of the variation) discriminated between genomic variation in *H. japonica* and *O. femoralis*. A combination of MDS1 and MDS2 discriminated between the genomic variation in *N. ovocercia*, *S. japonicus*, *A. longispina*, *R. japonica*, and *I. nipponica*. Environmental variables, including altitude, water level, water temperature, and precipitation significantly affected the genomic variation in five of seven species (*N. ovocercia*, *H. japonica*, *R. japonica*, *O. femoralis*, and *A. longispina*) (*p* < 0.05). No correlation was observed between candidate SNPs and environmental variables for the remaining two species, *S. japonicus* and *I. nipponica*. Highest genomic variation was detected in Gifu region along a spatially structured environmental gradient, whereas the least was detected in Sendai region.

db-RDA analysis of allelic variation revealed 294 SNPs (6–92 SNPs per species) significantly associated with six environmental variables (*p* < 0.05; Table 2). Thirteen SNPs were correlated with more than one environmental variable. Overall, precipitation and altitude were the most frequently selected environmental variables by the db-RDA models (Table 2), and showed strong correlations with 65 and 47 SNPs, respectively, except in the case of *H. japonica* and *I. nipponica*. Among the seven species, *I. nipponica* displayed the highest number of correlations between environmental variables and SNPs.

### Comparative genome analysis

Multiple sequence alignments of candidate SNPs sequences using bowtie revealed an average of 98% nucleotide sequence similarity among the seven species. Mean nucleotide substitution rate within species per geographical region (Table 3) ranged from 0–0.137 (mean = 0.091), where *S. japonicus* had the largest intraspecific genetic variation throughout the four studied regions (overall mean for four regions= 0.137). Six of seven species showed the highest mean nucleotide substitution rate in Gifu region (Table 3). Mean nucleotide substitution rate within geographical regions (range = 0.046–0.160, mean = 0.089) for seven species was significantly lower than that between geographical regions (range = 0.094–0.300, mean = 0.19) (*p* < 0.05). The Gifu region showed the highest mean nucleotide substitution rate between species (0.16) compared with other regions (range = 0.046–0.075).

**Table 3.** Mean nucleotide substitution rate within seven species distributed across four regions of Japan based on 4,251 candidate SNPs. Numbers in bold indicate the highest values

| Species | Matsuyama | Gifu | Sendai | Sapporo |
| --- | --- | --- | --- | --- |
| *Nemoura ovocercia* | 0.015 | **0.058** | 0.018 | 0.069 |
| *Haploperla japonica* | 0 | **0.317** | 0 | 0 |
| *Stavsolus japonicus* | **0.315** | 0.092 | 0.058 | 0.197 |
| *Rhabdiopteryx japonica* | 0.028 | **0.247** | 0.009 | 0.008 |
| *Obipteryx femoralis* | 0.018 | **0.115** | 0.026 | 0.004 |
| *Isoperla nipponica* | 0.116 | **0.301** | 0 | 0.259 |
| *Amphinemura longispina* | 0.035 | **0.213** | 0.037 | 0.013 |

DNA sequences of 4,251 candidate SNPs were subjected to tBLASTx search using the NCBI database in order to identify their gene functions. Of these, 3,868 sequences (91% sequences of the candidate SNPs, 4% of the total loci) were found to have matched with functional gene regions of other organisms (such as Drosophila and Lepidoptera, >90% similarity) based on Gene Ontology analysis using the UNIPROT database. Functions of most of these genes include metabolism and magnesium and citrate transport in the cuticle. However, confirming the expression of these functions in the insect genome requires additional examination along with other approaches such as gene expression analysis and proteomics.

We identified an average of 205 SNPs unique to each geographical region (range = 1–98 SNPs per species per region) (Table 4). Across all species, the highest number of unique SNPs occurred in Gifu (133 SNPs) and the lowest number in Sendai (10 SNPs). Among the seven species, *R. japonica* in Gifu showed the highest number of unique SNPs (98 SNPs) (Table 4). Unique SNPs showed high mean pairwise nucleotide substitution rate within species ranging from 0.223 in *H. japonica* to 1.012 in *S. japonicus*.

**Table 4.** Numbers of unique SNPs to each geographical region per species and mean nucleotide substitution rate of the unique SNPs within species

| Species | Matsuyama | Gifu | Sendai | Sapporo | Mean nucleotide |
| --- | --- | --- | --- | --- | --- |
|  |  |  |  |  | substitution rate |
|  |  |  |  |  | within species |
| *Nemoura ovocercia* | 6 | 6 | 4 | 14 | 0.407 |
| *Haploperla japonica* | 0 | 8 | 0 | 0 | 0.223 |
| *Stavsolus japonicus* | 8 | 6 | 0 | 2 | 1.012 |
| *Rhabdiopteryx japonica* | 6 | 98 | 2 | 7 | 0.331 |
| *Obipteryx femoralis* | 0 | 5 | 0 | 3 | 0.363 |
| *Isoperla nipponica* | 1 | 6 | 0 | 1 | 0.903 |
| *Amphinemura longispina* | 4 | 9 | 4 | 5 | 0.511 |

## Discussion

Genome-wide RAD-Seq analysis of seven stream stonefly species conducted in this study successfully identified candidate SNPs associated with environmental conditions along a nationwide latitudinal gradient in Japan. A total of 4,251 candidate SNPs in the stonefly genome were potentially associated with local environmental variations, of which 294 SNPs were significantly associated with six environmental variables and 205 were unique for a particular geographical region.

A substantial proportion (>70%) of genomic variation at the candidate SNPs was shaped by environment conditions for five of the seven species of stoneflies. Precipitation and altitude were most commonly correlated with allelic variation at candidate SNPs. Precipitation in the Japanese archipelago is abundant throughout the islands all year round (56). Regions in southern Japan, such as Matsuyama, experience the highest amount of rainfall in summer, whereas those in northern Japan, such as Sendai and Sapporo, experience homogenous rainfall throughout the year (Japan Meteorological Agency, www.jma.go.jp). Precipitation has been acknowledged as one of the principal environmental variables that shape aquatic insect distribution and diversity (57). Precipitation is also known to influence adaptive genetic divergence among populations in a variety of plants (e.g. *Sorghum bicolor*, 5) based on SNPs analysis. Our results showed that precipitation explained a substantial proportion of SNP variation, which was likely associated with the local adaptations of stonefly populations.

In addition to precipitation, altitude of the geographical regions was also associated with the genomic differentiation of stonefly species. Altitude is a long-term studied environmental variable implicated in the distribution (24) and genetic differentiation (58) of stoneflies species. In this study, sampling sites within a geographical region were located at different altitudes (Additional file 1: Table S1). It is, therefore, possible that population differentiation and local adaptation of stonefly species were affected by altitude variation within a region. Similar results have been commonly observed in aquatic insects based on mitochondrial DNA sequence variation (59), and in other taxa, including anemonefish (15) and sorghum (*Sorghum bicolor*; 14) using SNP genotyping. Together these data suggest that environmental requirements of stoneflies species play an essential role in their local adaptation and consequently, in adaptive genetic variation.

Among the four sampling regions of Japan, Gifu demonstrated the highest intraspecific genomic variation of candidate SNPs for six of seven species, and harbored 67% of total unique SNPs observed. This suggests that Gifu harbors a high level of genetic diversity. Geographically, Gifu is located in the middle of Japan surrounded by the Japanese Alps. This region has high insect diversity because of the geological formation history of the Japanese islands (56, 57). Ancestral Japanese landmasses were located along the borders of two major tectonic plates, namely the Eurasian (south Japanese landmass) and North American (north Japanese landmass) plate (60). Both landmasses had independent geological histories until their union approximately 20 million years ago (61) and the consequent formation of the Japanese Alps. The formation of the Japanese Alps has greatly contributed to the evolutionary process (species diversification) of Japanese aquatic insects (56) and the proteome variations of stoneflies species (62). However, no prior studies explored the geographic effect on intra-specific evolutionary patterns. Our findings provide an invaluable resource for the identification of genes associated with diverse traits of environment importance. Natural SNP variation influenced by environmental conditions has helped to detect genes associated with domestication (63) and climate change (64). Thus, multiple-species studies at the molecular level have the potential to detect hotspots of genetic diversity, thereby having implications for future research on river management and climate change. Moreover, multiple-species approaches may be useful in understanding how environmental pressure acts to shape genomic variation of a community in a region rather than that of an isolated species.

Genome–genome comparison of the candidate loci among the seven species showed high genomic similarity (98% nucleotide sequence similarity) and low mean nucleotide substitution rate (0.089) within their geographical regions. An evolutionary process that could lead to similar genomic profiles as a consequence of environmental conditions is parallel evolution (65). Parallel evolution occurs when phenotypes evolve in the same way in different species descending from the same ancestor because of their adaptation to similar habitats, and is a strong evidence of natural selection in adaptive evolution (66, 67, 68). Parallel divergence could reflect similar selective pressures acting on regions in different localities, geographically non-associated genetic traits, and dependent genetic drift (66, 68). Previous studies have demonstrated that similar environmental pressure can lead to parallel evolution of the genes under selection. For example, similar pattern of local adaptation of two stick insect populations under similar conditions of nutritional balance (*Timema cristinae*; 66), similar expression patterns of latitude-driven genes (as cytochrome b) in two *Drosophila* species (*D*. *melanogaster* and *D*. *hydei*; 67), and genes associated with morphological traits (as body shape, etc.) in two populations of three-spined stickleback (*Gasterosteus aculeatus*; 68) were reported. It is considered that mutations and evolutionary forces, as stabilizing selections that are invoked to maintain the genetic polymorphism among populations, will only play a role in maintaining genetic stability in species in a stable environmental over long periods of time (69).

## Conclusions

Our study demonstrated that local environmental conditions and ecological requirements of organisms play key roles in their evolutionary adaptation. Given the advantage of RAD-Seq, as finding similar genome regions (34, 35) and the repeatability of the experiments on multiple-species, we could affirm that our approach could detect signals of genome variation throughout local adaptation and could be used in future studies. To further understand the influence of environmental conditions in the genetic variation of locally adapted species, studies incorporating a larger number of samples and /or environmental variables combined with gene expression analysis should be conducted.

## Abbreviations

SNPs: single nucleotide polymorphism
PCoA: principal coordinate analysis
LFMM: latent factor mixed model
db-RDA: distance-based multivariate redundancy analysis
GO: gene ontology

## Acknowledgements

For logistic support in the fieldwork, we kindly thank Sueyoshi Masanao, Terutaka Mori, Ryota Kawanishi, and Kei Nukazawa. We also acknowledge Hiroki Hata for the assistant with RAD-seq analysis. Finally, we are grateful to Joeselle Serrana and anonymous reviewers for constructive comments on an earlier version of the manuscript.

## Availability of data and materials

The following data will be deposited at Dryad up to journal acceptance: structure files, candidate SNPs obtained using the LFMM function in R program and R code used in this analysis. The VCF files of SNP genotype output for stack pipelines for seven stoneflies species were submitted to European Variation Archive (EVA).

## Author’s contributions

M.G. and K.W. designed the study. Sampling, laboratory analyses, and bioinformatics analyses were conducted by M.G. All the authors contributed to the writing of the manuscript.

## Consent for publication

No applicable.

## Ethics approval and consent to participate

No applicable

## Additional files

**Additional file 1: Table S1**. Sampling sites and associated meteorological data

**Additional file 2: Table S2**. Number of individuals per sampling site. Geographical Regions: M, Matsuyama; G, Gifu; S, Sendai; and Sa, Sapporo. Sampling site codes are shown as described in additional file 1

**Additional file 3**: **Appendix S1.** *de novo* assembly filtering and post-assembly filtering

**Additional file 4: Table S3**. Number of loci by different parameter combinations evaluated for the *de novo* assembly

**Additional file 5: Table S4**. Number of loci after final data filtering using the population program of the STACKS program

**Additional file 6: Table S5**. F_*st*_values after final data filtering using the population program of the STACKS program

**Additional file 7: Table S6.** Numbers reads per species per region using the parameter combination m3 M2 n2 for the *de novo* assembly after the population program by STACKS

**Additional file 8: Table S7.** Numbers of loci per species per region using the parameter combination m3 M2 n2 for *de novo* assembly after population program by STACKS

**Additional file 9: Figure S1.** Flow chart of data filtering steps

